# Regulation of DNA (de)methylation positively impacts seed germination during seed development under heat stress

**DOI:** 10.1101/2021.02.25.432822

**Authors:** Jaiana Malabarba, David Windels, Wenjia Xu, Jerome Verdier

## Abstract

Seed development needs the coordination of multiple molecular mechanisms to promote correct tissue development, seed filling and the acquisition of germination capacity, desiccation tolerance, longevity and dormancy. Heat stress can negatively impact these processes and upon the increasing of global mean temperatures, global food security is threatened. Here, we explored the impact of heat stress on seed physiology, morphology, gene expression and methylation on three stages of seed development. Notably, Arabidopsis Col-0 plants under heat stress presented a decrease in germination capacity and also a decrease in longevity. We observed that upon mild stress, gene expression and DNA methylation were moderately affected. Nevertheless, upon severe heat stress during seed development, gene expression was intensively modified, promoting heat stress response mechanisms, including the activation of ABA pathway. By analyzing candidate epigenetic marks using mutants’ physiological assays, we observed that the lack of DNA demethylation by *ROS1* gene impaired seed germination by affecting germination-related genes expression. On the other hand, we also observed that upon severe stress, a large proportion of differentially methylated regions (DMRs) were located in promoters and gene sequences of germination-related genes. To conclude, our results indicate that DNA (de)methylation could be a key regulatory process to ensure proper seed germination of seeds produced under heat stress.

**Graphic Abstract:** 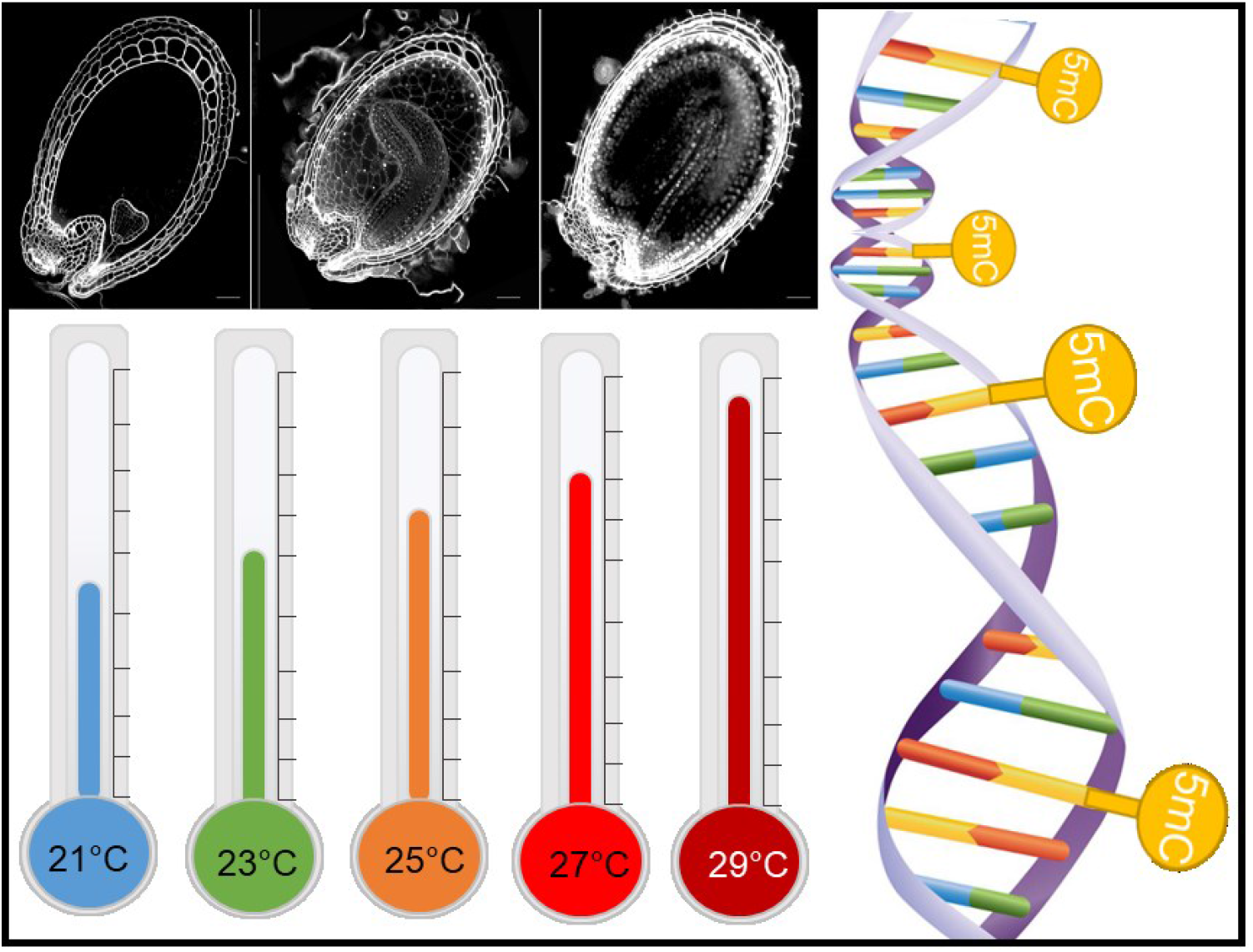

## Introduction

The accumulative anthropo-morphic greenhouse gas emissions causing climate change have a diverse and mainly negative impact on life on earth. The mean temperatures are increasing worldwide (0.35°C from 1979-2003) and also the range of diurnal temperature is higher since the minimum temperature increases much faster than the maximum temperature rate (1.13°C from 1979-2003) [1–3]. Even seasons are changing, with earlier biological spring and later biological winter [4]. The effect of global warming induced-heat stress on plants is strong and diverse, such as the advance in flowering time [5–7], modifications in plant architecture [8–10], decrease in grain yield [11], decreased in seed dormancy [12,13], the shift of plant establishment in higher altitudes [14], decrease of crops production such as maize and wheat [15], all of those posing a great threat to agricultural safety and ecological diversity. Concerning plant life cycle transitions, both timing of germination and reproduction are determined by temperature [16,17]. Therefore, the control of important seed traits has the potential to be seriously impacted by climate change [18]. Studies showed that heat stress can impact seed development and reduce seed size [19,20]. Along with physiological observations, plants’ molecular responses to heat stress has been extensively studied in the past decades. Heat stress factors (HSF) and Heat Shock proteins (HSP), calcium channels, phytohormones, chaperones, and other secondary metabolites have an important role in heat stress responses [21,22]. Moreover, a transcriptional reprogramming enables plants to cope with heat stress, by upregulating specific proteins, as kinases and transcription factors, along with direct stress protection genes, as detoxifying enzymes and osmoprotectants. On the contrary, growth-related genes are generally repressed [23]. Gene transcription is connected to its epigenetic state, therefore gene expression regulation is based on chromatin modifications (e.g. histone acetylation, methylation, phosphorylation and ubiquitylation) and DNA modifications (e.g. cytosine methylation) [24,25]. These modifications can be triggered by developmental or environmental factors and by consequence, they modulate chromatin architecture without changing the genomic sequences, but only the accessibility of transcriptional machinery to specific genome regions [26,27].

However, the understanding of epigenetic regulation of heat stress response is still dawning. DNA methylation is the addition of a methylgroup to a DNA cytosine base, forming a 5-methylcytosine (5mC). This process occurs in three sequence contexts in plants: the symmetric CG and CHG contexts and the asymmetric CHH context, where H stands for A, T or C. DNA methylation can be divided into (i) *de novo* methylation, (ii) maintenance of methylation and (iii) demethylation processes, in which different proteins are involved. In plants, *de novo* methylation is catalyzed by DOMAINS REARRANGED METHYLTRANSFERASE 2 (DRM2), which is regulated by RNA-directed DNA methylation (RdDM) pathway [28]. On the RdDM pathway, the RNA polymerase IV (Pol IV) transcribes single-stranded RNA (ssRNAs) that are later synthesized into double-stranded RNA intermediates (dsRNAs) by RNA-DEPENDENT RNA POLYMERASE 2 (RDR2). These dsRNAs are precursors for RNase III-class DICER-LIKE 3 (DCL3) to process into 24-nt interfering RNAs (siRNAs) that will be incorporated into ARGONAUTE 4 (AGO4) to be base-paired with Pol V transcripts, resulting in the recruitment of DRM2 and DNA base methylation [28]. Maintenance of DNA methylation is dependent on the sequence context, as CG methylation is maintained by METHYLTRANSFERASE 1 (MET1) and DECREASE IN DNA METHYLATION 1 (DDM1), while CHG is maintained by CHROMOMETHYLASE 3 (CMT3), and CHH by DRM2 [29]. On the other hand, plant DNA demethylation is regulated by four bifunctional 5-methylcytosine glycosylases: REPRESSOR OF SILENCING 1 (ROS1), DEMETER (DME), DME-like 2 (DML2) and DML3, which remove the 5-methylcytosine from DNA through the base excision repair pathway [30]. DNA methylation regulates gene expression, normally by inhibiting transcription of genes and also maintains genome stability against transposable elements [29,31]. Heat stress can affect DNA methylation as it was demonstrated in *A. thaliana* leaves with the increase of global methylation and homologous recombination frequency [32]. The methylation processes protein, DRM2, NUCLEAR RNA POLYMERASE D1(NRPD1) and NRPE1 were also shown to be up-regulated upon heat stress and could contribute to increase DNA methylation as a heat stress response [33]. The RNA-dependent DNA methylation pathway (RdDM) is partially responsible for the transcriptional response to temperature stress [23]. Nevertheless, heat stress-induced global DNA methylation changes appears to be species and tissue-dependent. Global DNA methylation was found increased in *Quercus suber* grown at 55°C [34] and in *Brassica napus* [35] but decreased in *Gossypium hirsutum* [36] and *Oryza sativa* [20]. Furthermore, investigating heat stress impact on transgenerational memory showed phenotypic changes over generations [32,37–39]. Heat stress ancestral exposure over two consecutive generations increased fitness in the F3 heat-treated Arabidopsis plants [40]. This epigenetic inheritance could promote genomic flexibility and adaptive advantages for plant fitness under heat stress conditions [41].

Thus far, the majority of heat stress studies focused on vegetative tissues with short-term heat stress events. Given the importance of seeds for agriculture, we aimed to evaluate the impact of different heat stress intensities during the whole seed development in *A. thaliana*, as mimicking global warming conditions. In our study we evaluated the impact of mild and severe heat stress on embryo growth and morphology; seed germination and longevity; gene expression and DNA methylation.

## Material and Methods

### Biological Material and Heat Treatment

*Arabidopsis thaliana* wild-types Columbia-0 (Col-0) and Wassilewskija (WS); and homozygous insertional mutants (T-DNA) *cmt3* (in WS background) [42] and *ros1-4* (SALK_135293 in Col-0 background) [43] were used in this work. Plants were grown in 6×6 cm square pots filled to the top with Tray substrate (Klasmann-Deilmann, Bourgoin Jallieu, France). Plants were kept at long-day conditions in a growth chamber with controlled humidity (60–70%) and average temperature of 23°C (24°C/22°C day/night) with a 16h-light/8h-dark photoperiod with a light intensity of 150 mmol.m^2^.s^-1^. Plants received watering 3 days per weeks on the base of their pots, for capillarity absorption by the substrate.

For heat stress treatment during seed development, as soon as the first flower buds appeared on the base of the leaf rosette, plants were transferred to growth chambers with the same light, photoperiod, humidity and watering conditions, but with different temperature ranges: 21°C (22°C/20°C day/night), 25°C (26°C/24°C day/night) or 27°C (28°C/26°C day/night) or 29°C (30°C/28°C day/night). A parallel set of control plants were continually grown in the 23°C growth.

### Physiological and phenotypic characterizations

Fully matured seeds from all genotypes were equilibrated at 44% of relative humidity (RH), using a saturated solution of K2CO3 at 20°C for 3 days, and then used for the physiological analysis. Col-0 plants at 29°C heat stress treatment produced insufficient seed number for all the experiments to be performed.

For water content analysis, seeds were samples at 4, 6, 8, 10, 12 and 14 DAF (days after fertilization). Three replicates of 50 seeds were used for determination of fresh weight and dry weight after 2 days drying at 96°C.

Protein content was characterized using a CHNS elemental analyser, which measured the percentage (w/w) of nitrogen using flash combustion of the sample based on the “Dumas” method. Mature seeds were ground in liquid nitrogen and dried in an oven at 90°C for 48 hours. Then, triplicates of approximately 5 mg of powder were analyzed using an Elementar Vario Micro cube analyzer (Germany). Protein content was estimated from the nitrogen content using the Jones Factor[44], which is used to convert the nitrogen concentration into total protein content.

The germination capacity was accessed by germinating triplicates of 100 dried seeds on Whatman paper No1 imbibed with 1ml of autoclaved water in a 3 cm diameter Petri dishes at 20°C under a 16h/8h photoperiod for eight days. Germination was scored every 2 days as seeds exhibiting a protruded radicle.

Seed longevity was evaluated by using accelerated (or artificial) ageing. Seeds were incubated in high humidity chambers (75% of RH from a saturated solution of NaCl) at 35°C in dark. At various intervals of storage, 100 seeds were retrieved and imbibed with 1ml of autoclaved water for germination assay to calculate the survival percentage of aged seed lots. Then, we modeled the seed longevity curve, characterized by the loss of seed viability along different ageing time. A longevity indicator was assessed as the storage time required for the seed batch to lose 50% of germination and called P50. Two-way ANOVA followed by Tukey’s multiple comparison test was used to determine the statistical significance of different time-points and/or genotypes. The exact statistical analysis parameters used are described in each related figure legend. Graphical creations and statistical analyses were performed using Prism 8.0.1 (GraphPad).

### Morpho-anatomical measurements

For evaluation of embryo development at different temperatures, Col-0 seeds were collected at different embryo stages: globular, transition, heart, torpedo, walking stick and mature. Seeds were cleared using the pseudo-Schiff propidium iodide staining as detailed [45].

Mature and dried seeds were used to measure seed width and length. SEM (scanning electron microscopy) pictures of seeds were acquired with the Phenom Pro Desktop SEM (Phenom-World) and all the samples were directly imaged, without sample preparation nor modification. For all measurements, seeds were obtained from 4 plants per replicate, and three replicates from three independent experiments were used for these measurements. Two-way ANOVA followed by Tukey’s multiple comparison test was used to determine the statistical significance of different time-points and/or genotypes. The exact statistical analysis parameters used are described in each related figure legend. Graphical creations and statistical analysis were performed using Prism 8.0.1 (GraphPad).

For evaluation of embryo survival, mature and dried seeds were incubated in a using 1% of 2, 3, 5 triphenyl tetrazolium chloride (Tetrazolium Red-TTZ-Merck, Kenilworth, NJ, USA) in a 50 mM phosphate buffer (pH 7.0) for 24 h at 30°C as previously described [19]. *ros1-4* mutant seeds were separated between normal-shaped and wrinkled-shaped seeds. After 24h, seeds were crushed between two glass laminas for embryo release to get in contact with the TTZ solution. Samples were then incubated at 25°C for two hours before macroscopic observation and imagery at stereo microscope Olympus SZX16.

### Total RNA Isolation and Transcriptomic Sequencing

Col-0 seeds were collected at embryo stages: heart, bent and mature, and developed at 23°C, 25°C and 27°C temperatures. *ros1-4* seeds were sampled at the mature stage and germination stages II and IV [46]. All samples were ground with micro-pistons and liquid nitrogen. Seed powders at germination stages II and IV were incubated in NucleoSpin® RNA Plant and Fungi kit lysis buffer with 1% of polyvinylpyrrolidone (PVP-40), followed by incubation at room temperature for 10 minutes and centrifugation at 8000G for 5 minutes before transferring supernatant to extract RNA, as described previously [47]. Total RNA was extracted from 100 freshly harvested seeds in three biological replicates using the NucleoSpin® RNA Plant and Fungi kit (Macherey-Nagel, Düren, Germany), according to the manufacturer instructions. RNA quantity and quality were measured using a NanoDrop ND-1000 (NanoDrop Technologies). cDNA library preparation and single-end sequencing (SE50, 20M) were outsourced to the Beijing Genomics Institute (BGI, https://www.bgi.com) using the DNBseq sequencing technology. After quality control, high quality reads were mapped on Arabidopsis reference transcriptome version 11 (Araport11) using quasi-mapping alignment and quantification methods of Salmon algorithm v.1.2 [48]. For gene expression analysis, raw RNA-Seq data were first normalized as Transcripts Per Kilobase Million (TPM). Transcripts with an average above 0 TPM in at least one developmental stage/tissue and with a coefficient of variation of log2 TPM > 0.05 among all developmental stages were retained for further analysis, resulting in 27,587 genes. Differentially expressed genes (DEGs) were determined using DESeq2 package (v1.22.2) [49], in which genes with log2 FC >0 or <0 and p-adjusted value of <0.05 for multiple testing with the Benjamini-Hochberg procedure which controls false discovery rate (FDR) were considered as differentially expressed. Gene annotation and GO terms were assigned according to the Araport11 annotation version of Arabidopsis genome. Over representation analysis (ORA) used GO enrichment terms were performed using Clusterprofiler [50] package in RStudio (version 1.3.1073) applying adjusted p-value cut-off <0.5 obtained from Bonferroni procedure.

Data mining for transcriptomics of Col-0 seeds from fresh seeds to seed imbibition for 48h was obtained from Gene Expression Omnibus (GEO, GSE94459) as publicly available dataset [51]. Raw data were mapped/quantified using Salmon with the same parameters described above. ImpulseDE2 algorithm [52], available in R, was used to unravel differentially expressed genes along the germination phase using the ‘case-only’ differential expression analysis and a adjusted p-value cutoff of 1%.

### DNA Isolation and Whole-Genome Bisulfite Sequencing

Col-0 seeds were collected at embryo stages bent and mature, and developed at 23°C, 25°C and 27°C average temperature. Two biological replicates were used for each stage and condition. Genomic DNA was extracted from the samples using the NucleoSpin® DNA Food kit (Macherey-Nagel, Düren, Germany), according to the manufacturer instructions, then quantified using a NanoDrop ND-1000 (NanoDrop Technologies). DNA samples were sent to to the BGI and library construction, bisulfite treatment using a ZYMO EZ DNA Methylation-Gold kit and paired-end sequencing using an Illumina Hiseq 2500 (PE100 20M) were outsourced to this company. Sequencing quality was verified by FastQC and clean reads were subsequently mapped to the *A. thaliana* reference genome index sequence version 10 (TAIR10) using Methylpy software [53]. Chloroplast genomic sequence from *A. thaliana* was used as an unmethylated control. After mapping, deduplication of sequences was performed and cytosine methylation sites were determined and quanitfied using Methylpy. Each context of methylation was considered independently: CG, CHG or CHH. Putative differentially-methylated regions (DMRs) were identified from merged differentially methylated sites (DMS) using Methylpy ‘DMRfind’ method.

## Results

### Impact of constant heat stress on *A. thaliana* seeds

To observe which range of constant heat stress would impact seeds and how this impact could influence seed development and important seed traits, such as germination capacity, longevity and seed filling, assays were conducted with Col-0 plants grown in optimal conditions in growth chambers until their reproductive stage (i.e. apparition of first flowers), then moved to growth chambers with an average temperatures of 21°C, 23°C, 25°C, 27°C and 29°C until seed maturity. Our focus on this study was the evaluation of seeds but we also observed that plants presented different phenotypes when grown at heat stress conditions. Plant biomass was increased, while the silique number and length decreased, as temperature raised (Supplementary Figure 1). The effect of heat stress in seeds is first observed at water content measurements from 4 DAF to 14 DAF. Seeds developed at 25°C presented lower water content than seeds at 21°C and 23°C, and seeds at 27°C presented decrease water content in all time points, from 4 DAF until 14 DAF. At 14 DAF, the water content of seeds at 27°C was 20% less than at normal temperature (Figure 1A). Heat stress also had a negative impact on protein content. Seeds from 25°C and 27°C presented decreased protein content when compared to seeds from 21°C and 23°C (Figure 1B). When evaluating germination capacity of mature seeds, 21°C and 23°C seeds presented no difference between each other, with almost 100% of seed germination. However, germination was negatively impacted by heat stress treatments. Seeds from 25°C, 27°C and 29°C presented progressively decreased seed germination, with −20% germination capacity at 25°C, −35% at 27°C and −45% at 29°C (Figure 1C). Seed longevity was accessed by artificial ageing and seed germination evaluation after successive storage time-points (6, 10, 14, 21, 28 days). The P50 for seeds from 21°C and 23°C seeds are of 14 days of storage. Nevertheless, for 25°C and 27°C, P50 was significantly decreased, with 50% of seeds incapable of germination after 10 days of storage (Figure 1D). We were also interested in evaluating the impact of constant heat stress on seed shape. By microscopically phenotyping seeds, we were able to observe the presence of wrinkled-shape seeds due to heat stress conditions. Seeds from plants at 23°C and 25°C presented mainly normal-shaped seeds, while seeds at 27°C presented normal and wrinkled seeds (17%) (Figure 1E). Concerning seed development, we did not observe any effect of heat stress on seed embryogenesis, since the anatomical evaluation of seed layers and embryo showed normal embryo development and growth (Supplementary Figure 2).

**Figure 1.**
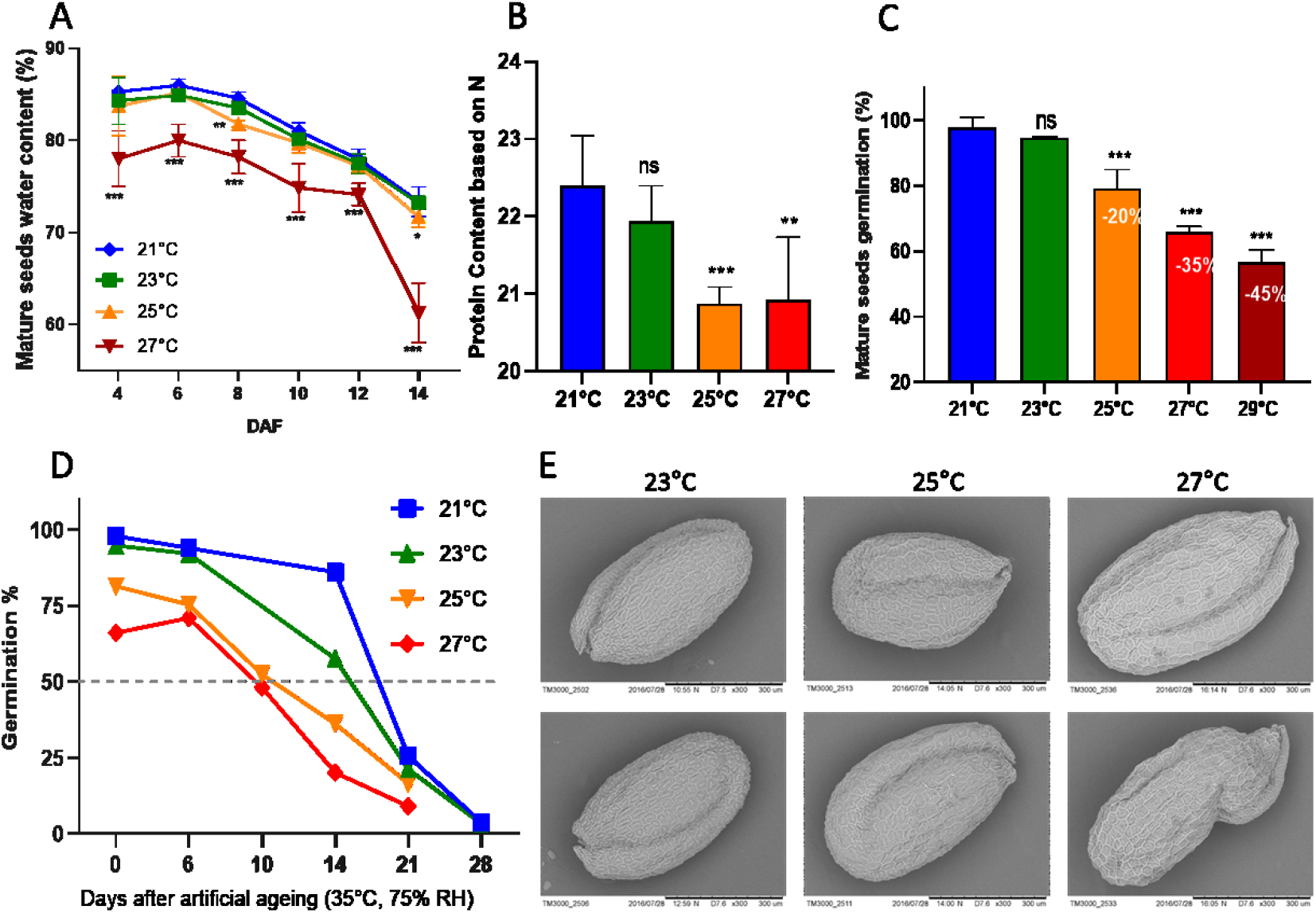
Physiological view of *A. thaliana* seeds under heat stress development. (A) Mature seed water content measured every 2 days for a duration of 14 days. Two-way ANOVA showing the difference between stress conditions and control conditions at each time point. (B) Protein content based on nitrogen content. Two-way ANOVA comparing 23°C, 25°C and 27°C against 21°C. (C) Percentage of fresh seed germination in which the numbers inside bars represent the decrease in seed germination upon heat stress conditions compared to control temperatures. Two-way ANOVA comparing 23°C, 25°C and 27°C against 21°C. (D) Seed longevity based on P50, in which artificial ageing was imposed for 6, 10, 14, 21 and 28 days. (E) Exemplary electronic microscopy of Col-0 seeds grown at 23°C, 25°C and 27°C. The below panel of 27°C condition shows a wrinkled-shape seed. Asterisks represent p-value significance: * p< 0.05; ** p< 0.001; *** p< 0.0001. Standard deviation is shown in A, B and C graphs.

### Gene expression during seed development under heat stress

Based on the physiological assays’ results, we defined three temperatures and three embryo developmental stages for further experiments. The stages of Heart (H), Bent (B) and Mature (M) embryo were sampled from plants grown at 23°C (control temperature), 25°C (Mild Stress, MS) and 27°C (Severe Stress, SS) (Figure 2A). We analyzed the dynamics of transcript abundance during seed development under control conditions and heat stress conditions by whole-transcriptome RNA-seq analysis. Differentially expressed genes (DEGs) were observed in each stage by comparing MS versus control and SS versus control condition. The amount of DEGs observed at 27°C was significantly higher than the amount of DEGs at 25°C. At heart stage, 27°C seed had 16-fold more up-regulated DEGs and 19-fold more down-regulated DEGs than at 25°C. The same happened at bent stage, with 31-fold more up-regulated DEGs and 22-fold more down-regulated than at MS condition, and at mature stage, seeds on severe stress presented 9-fold more up- and 7-fold more down-regulated genes than seeds on mild stress (Figure 2B). DEGs from severe stress were compared between stages and 533 genes were commonly up-regulated in all stages, representing around 19-25% from total DEGs of each stage, while 683 genes were commonly down-regulated in all stages analyzed, around 20-25% of total DEGs from different stages (Figure 2C).

**Figure 2.**
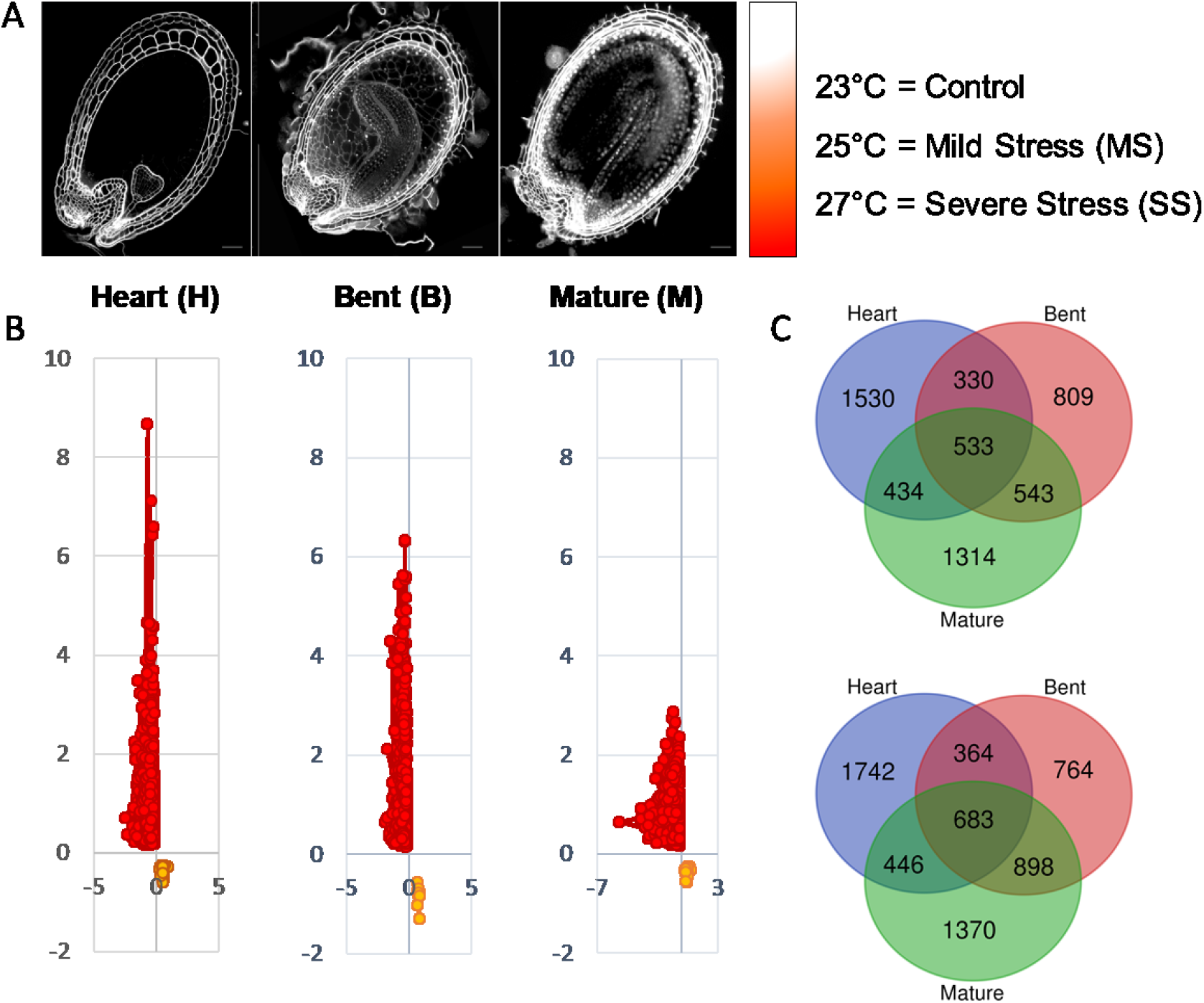
Transcriptomics analysis of Col-0 seeds developed under heat stress conditions. (A) RNA-seq samples were obtained during embryo developmental stages showed here in time-course order: Heart (H), Bent (B) and Mature (M). Average temperature of 23°C was used as control against Mild Stress (MS) at 25°C and against Severe Stress (SS) at 27°C. (B) Scatter plot showing the quantity of differentially expressed genes (DEGs) for MS in orange and SS in red. (C) Venn diagrams of DEGs from SS at each embryo developmental stage. Upper panels show up-regulated genes and lower panels show down-regulated genes.

We were interested in better understanding the function of the up- and down-regulated DEGs, during SS since few gene changes during MS, therefore we performed a gene set enrichment (GSEA) analysis using GO terms. The main biological functions of the up-regulated DEGs were response to abiotic stress such as heat stress, water deprivation, stomatal movement and response to hydrogen peroxide. Furthermore, secondary metabolism mechanisms were induced by severe heat stress, with the up-regulation of the phytohormones abscisic acid and jasmonic acid, as well as glucosinolates and suberin biosynthesis (Figure 3A). ABA signaling pathway was found up-regulated in all stages upon severe stress, while ‘positive regulation of germination’ terms were up-regulated only during heart stage (Figure 3A). Concerning the GO term enrichments of down-regulated DEGs, ribosome biogenesis and maintenance were present for all stages, along with the decrease of photosynthesis mechanism by repression of photosystems I and II, chlorophyll and thylakoids (Figure 3B). The over-representation analysis shows that the down-regulated DEGs are more homogeneous in their response to heat stress, but that the up-regulated DEGs present more distinct responses dependent on their development stage. Another point is that the specific DEGs of each stage, heart, bent and mature, showed a specific response at each stage against severe heat stress (Supplementary Figure 3). The complete differential expression analysis is available at Supplementary Table 1.

**Figure 3.**
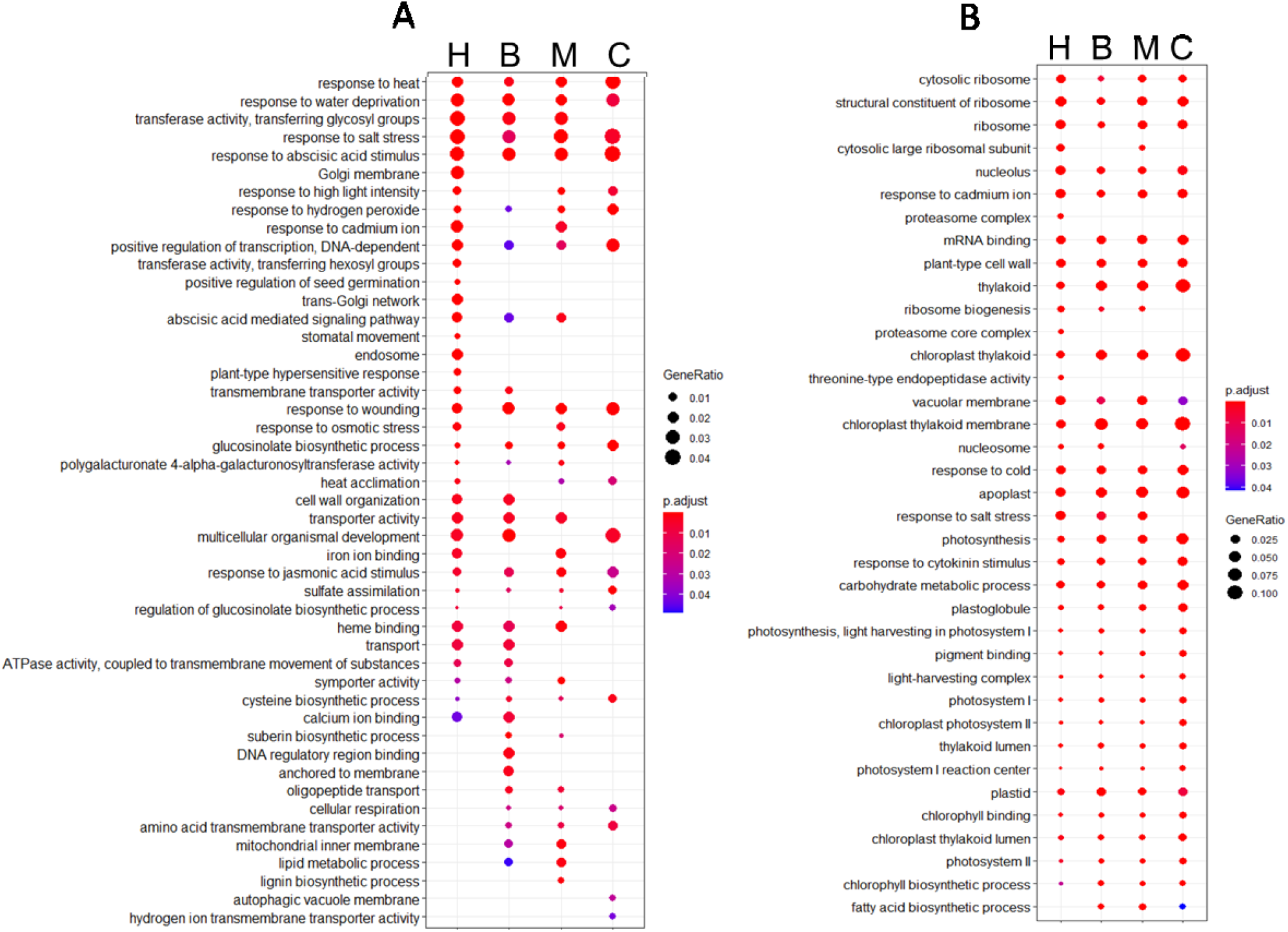
Gene set enrichment (GSEA) analysis of DEGs in Col-0 seeds under severe heat stress. (A) Functional enriched GO terms from up-regulated differentially expressed genes. (B) Functional enriched GO terms from down-regulated DEGs. The size of the dot represents the gene count. The totality of differentially up- or down-regulated genes in each stage was used to perform a hypergeometric test, and the p-values were converted to false discovery rate (FDR)-corrected p-value as shown in colours, the red colour being more significant than the blue colour. H= heart, B= bent, M= mature and C= common to all stages.

### Severe heat stress effect on methylation-related mutants

Our transcriptomic results also showed differences in the expression of genes related to chromatin organization and DNA methylation on seeds upon heat stress. The expression of *CMT3* and *ROS1* genes were affected by mild and severe heat stress in the embryo stages analyzed (Figure 4A). *CMT3* expression increased at heart stage at 27°C, and at bent stage, its expression increased at mild and severe stress. However, at mature stage, *CMT3* gene expression remained stable upon mild and severe stress. For the demethylase gene *ROS1,* severe heat stress dramatically impacted expression at bent and mature stages (Figure 4A). *ROS1* was also differentially expressed (down-regulated) in all developmental stages upon severe stress. Therefore, to tackle the role of these genes on seeds under severe heat stress development we decided to further investigate both gene mutants. We produced seeds from *cmt3* and *ros1-4* mutant plants grown at 23°C and 27°C for anatomical and physiological assays. First, we evaluated their seed gemination rate. At 23°C, mutants *cmt3* and *ros1-4* seeds presented no difference from its background, WS and Col-0, respectively. Upon severe heat stress, seed germination was not affected for both WS and *cmt3* genotypes. However, for Col-0 and *ros1-4*, severe heat stress decreased seed germination. Col-0 germination decreased in half while on the loss of function mutant *ros1-4*, severe heat stress caused an almost complete loss of germination capacity (Figure 4B). To better understand the consequences of severe heat stress on these mutants’ seeds, we observed and measured seeds with electronic microscopy (Figure 4C ad 4D). From all the genotypes analyses, *ros1-4* under 27°C presented the most drastic phenotype. 86% of *ros1-4* seeds were wrinkled and deformed, some with the aspect of ‘empty seeds’ (Figure 4C and 4D). Concerning seed measurements, there was no significant difference in length and width for *ros1-4* at 27°C, nonetheless, there was a significant difference between *cmt3* length and width at 23°C when compared to its background, as well as 7% of its seeds produced at 27°C displayed the wrinkled phenotype (Figure 4D). By observing *ros1-4* seed phenotype upon severe heat stress, we raised the hypothesis that *ros1-4* embryos were defected or lost their viability. To observe embryo survival, we used Tetrazolium Red to Formazan assay, in which living tissue or cells containing dehydrogenase enzymes thanks to hydrogen released during cell respiration, will reduce tetrazolium chloride to formazan, a reddish, water-insoluble compound [54]. Therefore, living cells will be red-stained indicating normal cell respiration. *cmt3* displayed red-coloured embryos at control and upon severe heat stress. *ros1-4* seeds at 23°C also displayed normal anatomic and reddish embryos. Under 27°C, *ros1-4* seeds were divided between normal and wrinkled-shape for preventing erroneous interpretation of results. Normal shape seeds presented viable embryos as well as non-viable embryos, while the majority of wrinkled seeds were empty or presented embryos with arrested development, as seen in Figure 4C.

**Figure 4.**
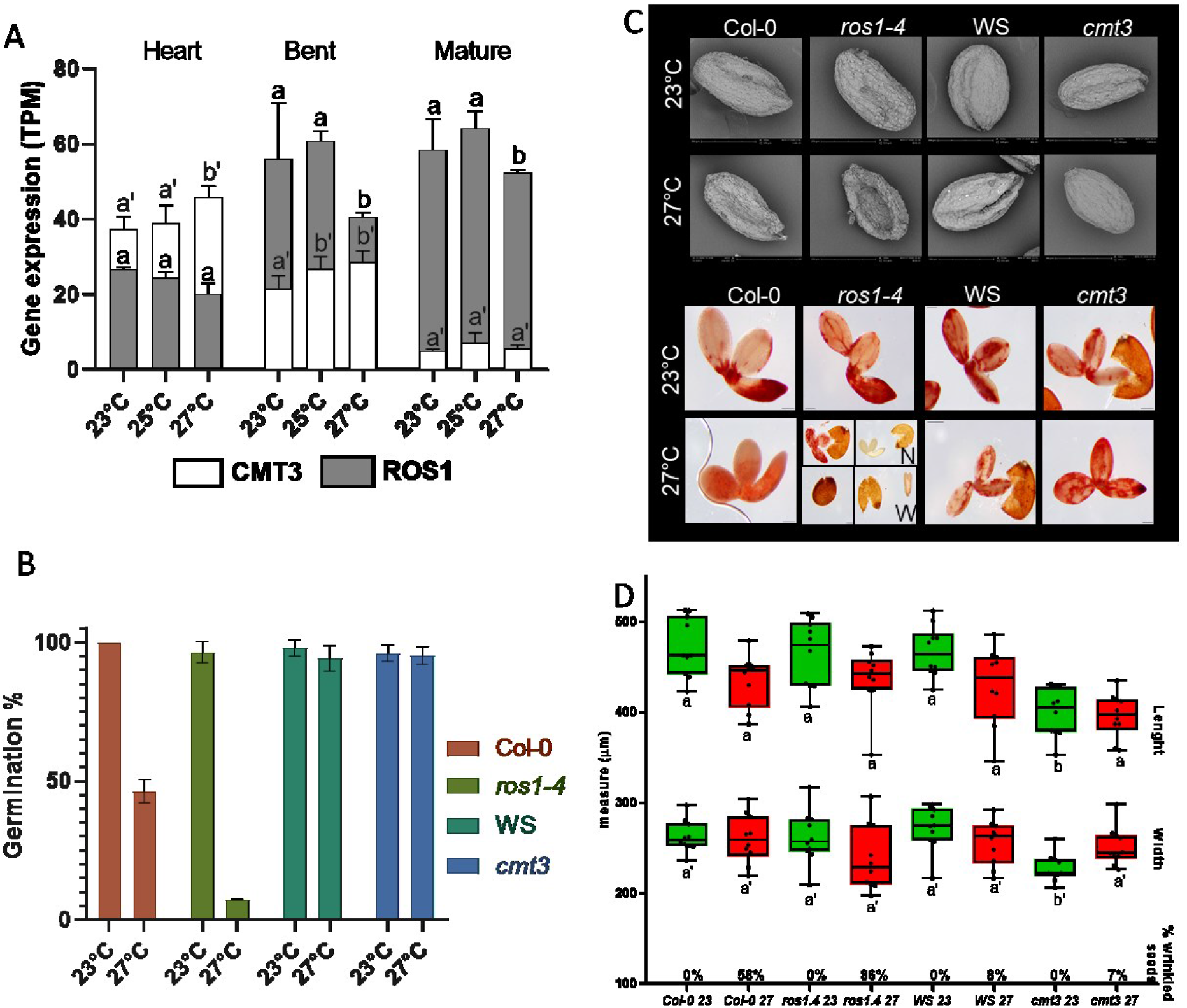
Evaluation of *CMT3* and *ROS1* expression and physiology of *cmt3* and *ros1-4* mutants. (A) Gene expression in TPM of *CMT3* and ROS1 at heart, bent and mature stages of embryo development during control temperature (23°C), MS (25°C) and SS (27°C). *CMT3* is represented by white bars while *ROS1* is represented by grey bars. Standard deviation is shown. Two-way ANOVA comparing differences in gene expression at 23°C, 25°C and 27°C for each genotype at same stage. Letters represent statistically significant differences with a p-value <0.001. Letters a or b used for *ROS1* and letters a’ or b’ used for *CMT3.* (B) Seed germination for wild-type (Col-0 or WS) and mutants *(ros1-4* or *cmt3)* genotypes at 23°C (green) and 27°C (red) average temperature. (C) Panels show embryos and seed coats from wild-types (Col-0 or WS) and mutants *(ros1-4* or *cmt3)* genotypes at 23°C and 27°C average temperature. The upper panel is composed of electronic microscopy photos for phenotype observations and lower panel shows embryos survival evaluation with TTZ. *ros1-4* at 27°C was divided into normal-shaped seeds (N) or wrinkled-shaped seeds (W). (D) Seed measurements of length and width in micrometers. Boxes and whiskers showing minimum and maximum with average line from all observations. Green bars represent temperature 23°C and red bars represent the temperature at 27°C (SS). Two-way ANOVA comparing differences size between wild-types (Col-0 or WS) and mutants *(ros1-4* or *cmt3*) respectively, at 23°C wild-type against 23°C mutant, and 27°C wild-type against 27°C mutant. Letters represent statistically significant differences with a p-value <0.001. Letters a or b used for length measures and letters a’ or b’ used for width measures. Percentage of wrinkle-shaped seeds for each genotype at 23°C and 27°C is shown at the base of the x-axis.

### Whole-genome DNA methylation of seeds under heat stress

To understand the precise role of DNA methylation on seed response to heat stress, we performed bisulphite sequencing of the genomic DNA isolated from Col-0 seeds produced at 23°C, 25°C and 27°C average temperature, therefore control, mild and severe heat stress. We used the stages of bent and mature embryo for this analysis. About 20 million high-quality read pairs were generated for each sample and mapped uniquely to the Arabidopsis genome (TAIR10). The percentage of methylcytosines identified in each context did not change upon mild nor severe stress (Figure 5A). At bent stage, % of methylated cytosines without heat stress was of 19.7% for CG, 6.9% for CHG and 1.4% for CHH, while at mild heat stress was of CG=19.4, CHG=6.6 and CHH=1.3, and at severe stress CG=20.3, CHG=6.9 and CHH=1.4. The same tendency was observed at mature stage of seeds produced at 23°C (CG=19.3, CHG=7.5, CHH=2.1), 25°C (CG=20.1, CHG=7, CHH=2) and 27°C (CG=20.7, CHG=7.2, CHH=2.2) with a similar percentage of methylcytosines per context and per heat stress (Figure 5A). We identified differentially methylated regions (DMRs) upon mild heat stress and upon severe heat stress when comparing heat-stressed samples to control samples. DMRs were identified on gene sequences and 1Kb promoter gene regions. A total of 110 DMRs were identified, and from those, 87 unique genes were annotated (Figure 5B). Severe stress induced more DNA methylation changes than mild stress, with 33 DMRs at bent and 49 DMRs at mature stage, against 10 DMRs at bent and 18 DMRs at mature stage upon mild stress. Mature stage had a greater total amount of DMRs (67) when compared to bent stage (43). Interestingly, DMRs were also found in mitochondrial DNA, with 13 DMRs present upon mild stress and 1 DMR upon severe heat stress. From these, five DMRs were localized at mitochondrial coding or promoter gene regions. A complete list of the genes related to the DMRs and their contexts are on Supplementary Table 2.

**Figure 5.**
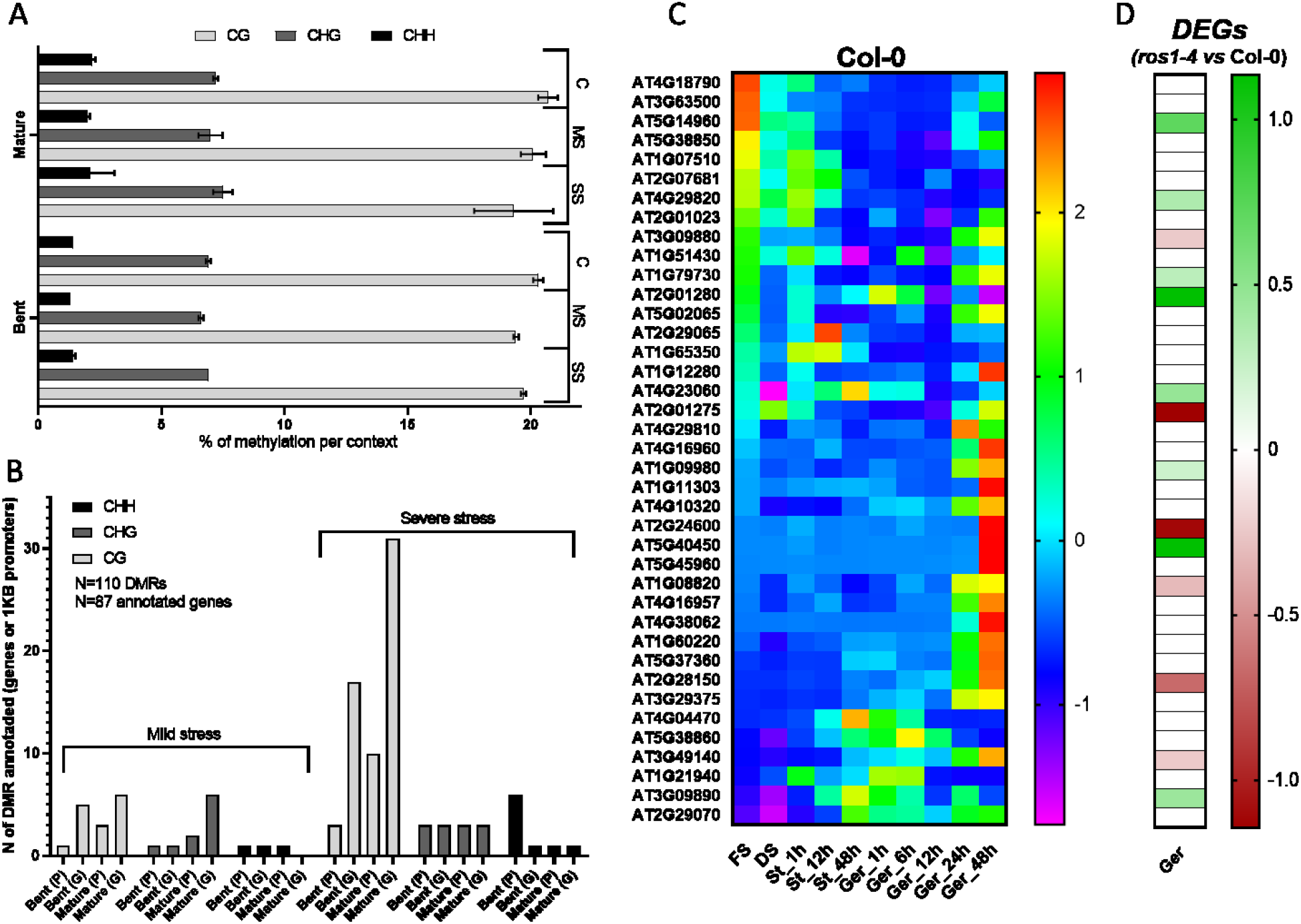
Methylome analysis and DMRs related to germination genes. (A) Percentage of cytosine methylation per context (CG, CHG, CHH) on bent and mature embryo stages of Col-0 seeds grown under Control temperature (C=23°C) or Mild Stress (MS=25°C) or Severe Stress (SS=27°C). (B) Amount of DMRs between control condition and MS or control condition and SS in all contexts (CH, CHG, CHH) for bent and mature stages at 1Kb promoter regions (P) and gene sequences (G). (C) Normalized expression (TPM) heat-map of 39 genes differentially expressed during seed germination and containing a DMR at mature stage with severe heat stress (MethDEGs). Expression in Col-0 seeds at conditions: fresh seed (FS); dried seed (DS); stratified and imbibed seeds at 4°C (St) for 1h, 12h and 48h; stratified and imbibed seeds at 20°C light for germination (Ger) for 1h, 6h, 12h, 24h and 48h. (D) Differential expression analysis of MethDEGs between *ros1-4* and Col-0 on three germination stages, whereas green indicates up-regulated genes and red indicates down-regulated genes in *ros1-4*.

Given that germination was affected by heat stress and that *ros1-4* presented a strong germination phenotype upon severe stress, we compared our DMRs annotated genes with germination-related genes identified by transcriptomics analysis. The result shows that 69% of the differentially methylated genes (39 out of 87 genes) were also differentially expressed genes during seed germination, therefore potentially involved in germination process. To examine the influence of differential methylation on differential gene expression, we analyzed expression profiles of these 39 methylated and differentially expressed genes (MethDEGs) between successive stages of seed development germination, from fresh seed, dry seed, stratified seeds at 4°C and seeds incubated at 20°C light for germination from 1h to 48h (Figure 5C). We observed two gene clusters: (i) genes with accumulated transcripts on mature seed, that are progressively decreasing in expression along with seed germination process (upper heat map), and (ii) genes that are induced upon stratification and seed imbibition, with an increased expression during seed germination with seed coat rupture and radicle expansion (middle-below heat map) (Figure 5C). We also performed RNA-seq with *ros1-4* seed grown at 23°C, using mature dried seeds, and two stages of germination: stage II (seed coat rupture and radicle point) and stage IV (seed coat rupture and radicle 0.5 mm expansion). 84% (32 genes) of the MethDEGs were found to be differentially expressed in *ros1-4* during germination when comparing mature stage of *ros1-4* against its samples at stage II or against stage IV of germination. We also performed a differential expression analysis between Col-0 and *ros1-4* (Figure 5D). On the *ros1-4* mutant, that shows decreased germination and decreased heat stress resistance, we observed that 14 MethDEGs were differentially expressed. From those genes, 8 were up-regulated and 6 were down-regulated in the mutant (Figure 5D). From the up-regulated genes, ELF7 (EARLY FLOWERING 7 - AT1G79730), DEL2 (DP-E2F-like transcription factor - AT5G14960) and BRF3 (TFIIIB-related factor - AT2G01280) genes are interesting because they were already shown to be required for seed dormancy, proper plant cell proliferation and elongation and thermotolerance. In detail, the ELF7 is required for the expression of the flowering repressor FLC [55] and is also part of the Polymerase II Associated Factor 1 Complex (PAF1C) which interacts with Reduced Dormancy 2 factor (RDO2). A studied with *elf7* mutant showed that seed dormancy was reduced, suggesting the role of this gene the control of seed germination [56]. DEL2 altered expression caused changes in cell division and elongation in the root meristem and affected the expression of several cell-cycle regulators [57]. Moreover, the BRF3 gene, which is part of the core unit of the RNA Pol III transcription complex [58], was shown to negatively regulates the thermotolerance in Arabidopsis [59]. The up-regulation of these genes in *ros1-4* during germination could be part of the answer for the decrease in *ros1-4* seed germination upon heat stress. The complete differential expression analysis between *ros1-4* and Col-0 is available at Supplementary Table 1.

## Discussion

### Constant heat stress does not affect seed embryogenesis but negatively impacts seed germination and longevity

Here we aimed to verify the impact of constant heat stress on seed development. To perform the experiments with *A. thaliana* we based our temperature selection on global warming projections. Initial projections estimated an increase of 1°C to 2°C on average global surface temperature, but recent projections foresee an increase by up to 4.8°C by the year 2100, depending on the level of greenhouse gas emission [60]. Along with this gloom data, climate change is projected to impose more extreme weather conditions, with the rise in global temperature including longer and hotter summers that possibly will severely disrupt plant growth and productivity [61].

We were keen to investigate if heat stress could disrupt seed embryogenesis, which is the developmental process that transforms a single totipotent cell, the zygote, into a mature embryo, by producing and developing the embryo first tissues precursors and first stem cells [62]. We observed that seed embryogenesis was not affected upon our heat stress assays. One explanation might be the strength of embryogenesis cell pattern in *A. thaliana.* Most flowering plants present apparently random and disorder embryogenic cell divisions while a few present a highly regular and predictable pattern, such is the case of the Brassicaceae family [63,64]. This is why most of the knowledge on plant embryogenesis comes from Arabidopsis research, again with the advantages of small genome size, rapid life cycle and amenability to genetic transformation [62]. Our assays showed that the majority of Col-0 seeds at severe stress still developed correctly and this fact could also be due to plants mechanism of basal thermotolerance, that is when plant survival above optimal temperatures is assured. Basal thermotolerance must be distinguished from acquired thermotolerance, that is survival to otherwise lethal heat stress, both being physiologically and molecularly different processes [65].

Even if we did not observe anatomical abnormalities during embryogenesis, we observed reduced silique number and length as temperature increased. As silique length is correlated with the number of seeds per silique [66,67], it is probable that less efficient fertilization lead to fewer seeds and shorter siliques. This was the case for heat stress assays in *A. thaliana* and in *Lycopersicon esculentum,* that negatively affected flower fertilization [67–70]. Flower infertility might also be produced when heat stress inhibits anther dehiscence, thereby reducing pollen release; and pollen viability by disrupting male meiosis [70–72]. Heat stress can also have an impact on the female gametes with malformation of the ovule and accessory tissues, as well as decrease of stigma receptivity [73,74]. In our assays, all plants were grown at control conditions until bolting, then plants were transferred to heat stress conditions as described. Nevertheless, as Arabidopsis flowers continuously appear, later-developed flowers might have been more stressed than first-developed flowers. This could be the explanation for shorter and fewer siliques, with less seeds.

Regarding heat stress induced impairment of seed essential traits, heat stress reduced germination by 30% to 50% and longevity by half. The negative impact of heat stress on germination has already been reported [75–77]. These major impact on seed germination and longevity can have disastrous consequences for plant species survival since these are key processes to ensure species propagation [78–80]. For promoting seed germination, two phytohormones are key players: Gibberellic acid (GA) and Abscisic acid (ABA) [81]. During seed germination ABA biosynthesis is repressed while GA increases [82]. The contrary is true to dormancy in which ABA content is high and GA is low [83]. We observed that upon severe heat stress, the ABA pathway was induced in all stages, and as the regulation of plant process upon abiotic stress have a modulated activity depending on the water status in the environment [82], the extremely low water content on seeds upon severe heat stress might have disrupted the regulation of germination process.

### DNA demethylation is partially responsible for ensuring seed germination under heat stress conditions

Our study had the objective of characterising the transcriptomic and epigenomic changes that occur on seeds when they are faced with a developmental environment with constant heat stress. We observed that the transcriptional answer to heat stress was significant when plants were grown on severe heat stress conditions and that DEGs upon severe heat stress from all the stages analysed were mainly related to abiotic stress response and the repression of photosynthesis machinery. These results reinforced previous works that showed that a myriad of cell processes is affected by heat stress, since even enzyme function can be disrupted by temperature and provoke metabolic imbalance [84,85], or even promote programmed cell death [86,87]. Moreover, a variety of membrane-linked processes are impacted due to changes in membrane fluidity and permeability [88,89]. By promoting membrane and protein damage, heat-induced oxidative stress is generated by the production of active oxygen species [90–93]. Taken together all the possible damaged, heat stress in plants is characterized by impaired translocation of assimilates and reduced carbon gain and reduced photosynthesis [90]. Heat stress influences plant growth, productivity and development, being more related to impaired development. But soon after the stress period, plants will reinitiate their developmental and growth program and end the transcriptional response to stress, that will provide successful recovery.

The epigenetic regulation of heat stress response varies, from DNA methylation [20,34,36], to histone modifications [20,34,36,94,95] and histone variants [96,97], and also microRNAs and siRNAs [39,98,99]. Nevertheless, these epigenetics responses are dependent on the heat intensity and duration, given that studies have been done with plants grown with heat stress or with peaks of heat stress with different durations and repetitions. To better understand the dynamics of DNA methylation on embryogenesis stages upon heat stress, we chose to examine closely two stages that represent the middle and the end of embryogenesis, bent and mature stages, respectively. When analysing the epigenetic changes on the DNA, we observed that global DNA methylation levels were not modified by heat stress. The Arabidopsis genome-wide methylation levels are of 24% on CG, 6.7% on CHG and 1.7% on CHH contexts [100]. We detected a slight difference from each context against data from leaves samples [100], but not between stages or upon mild or severe heat stress. This could indicate that different tissues have different DNA methylation levels and also that methylation is stably maintained thought-out the genome upon constant heat stress, like a basal DNA-methylation thermotolerance.

In plants, the DNA methylation homeostasis is regulated by DNA methylation and also by demethylation processes [27,30] in a way that active DNA methylation and demethylation are balanced in the cells [101]. Since we observed the differential expression of *CMT3* and *ROS1* upon heat stress, we aimed to investigate if maintenance of methylation by CMT3 and/or DNA demethylation by ROS1 could have a role on heat stress response on seeds. During the mature embryo stage, cell division is arrested, and therefore it suggests DNA replication and maintenance of DNA methylation by MET1 during replication is also stopped. Therefore, the creation of DMRs during this stage is hypothesized to be due to DNA demethylation processes [101]. Recent studies showed that ROS1 is active in developing seeds during late embryogenesis by promoting differentially methylated regions on seed-development-related genes, more specifically for the demethylation of endosperm-specific methylated regions [101,102]. At the mature embryo stage, we observed an increase in *ROS1* expression in normal conditions and a decrease in *CMT3* expression, however, severe heat stress repressed *ROS1* expression. Furthermore*, DML3* expression increased upon severe heat stress, while other demethylases, *DML2* and *DME* were repressed and methylases *DRM1, CMT2* decreased, and *RDM1, RDM4* and *CMT3* remain stably low. The same was detected by a recent study that showed that while *DME, DML2* and *DML3* were repressed, *ROS1* expression was induced at maturation stages [101]. In addition for *CMT3* expression, a study in rice seeds showed repression of its expression with a 48h of moderate heat stress (34°C) after fertilization, which was proposed to cause differences in the DNA methylation levels of *Fertilization-Independent Endosperm1* (*OsFIE1*), a member of Polycomb Repressive Complex2 (PRC2) [20].

After physiological analysis of both *cmt3* and *ros1-4* mutants, *ros1-4* presented an interesting phenotype, with seed germination dramatically affected by severe stress, and with deformed seeds and undeveloped embryos. The *ros1-4* mutant has a Col-0 ecotype background with T-DNA insertion in the *ROS1* gene, proved to cause complete loss of function of *ROS1* [103]. As *de novo* methylation is established through the RdDM pathway, *ROS1* antagonizes RdDM and RdDM-independent DNA methylation. This mechanism was shown to prevent the spreading of DNA methylation from transposable elements to protein-coding genes [104–106]. At the same time, *ROS1* expression is positively regulated by proximal RdDM-dependent TE methylation [107,108]. Seed establishment is largely influenced by the amount and diversity of compounds storage during seed maturation [109], whereas germination capacity is not completely dependent on seed reserve accumulation, since the germination of mutants deficient in lipid reserve mobilization pathways is only slightly affected [110–114]. Therefore, germination efficiency must be controlled by other yet unknown mechanisms, one on those being the DNA demethylation by ROS1. Here, we observed a set of germination-related genes that were differentially expressed upon severe heat stress and were also differentially methylated at seed maturity. When analyzing their expression pattern, we identified the remodelling of the transcriptome upon the steps culminating in germination. These changes were also observed in previous works that investigated the seed transition from fresh, dry seed stage, seed stratification, germination and post germination, following exposure to light [51]. Studies shown that during the seed maturation process a diverse set of molecules, such as proteins, lipids, sugars and transcripts will be accumulated to be used upon germination [113,115–117]. Upon seed imbibition, a metabolic switch happens and the metabolites accumulated during seed maturation are consumed, by being mobilized and/or degraded, also with that germination-associated gene expression programs start already during seed imbibition [80]. Thus, we can witness these two clusters of gene expressions in the MethDEGs germination-related as well.

Here we showed that mother-plants grown under heat-stressed generated seeds with DMRs located on germination-related genes. These DMRs could be conserved in the future generations, as many works presented the inheritance of epigenetic changes caused by heat stress by the next generations and that this transgenerational inheritance was maintained for at least three generations in a genotypic and phenotypic manner [32,37–39,95]. The epigenetic memory can increase plant fitness to provide better heat stress responses and therefore contribute to plant evolutionary adaptation [40,41]. Also, *ROS1* was shown to decrease its expression by heat stress and that this decrease was maintained in heat-stressed *A. thaliana* F1 descendants, which also presented earlier-bolting, as well as fewer and larger leaves [38].

Remarkably, five mitochondrial genes presented DMRs upon heat stress. These genes were all found to be highly expressed in seed development, imbibition and germination. Given that mitochondria are responsible for respiratory chain, these genes could be important actors during seed germination, and different DNA methylation status caused by heat stress could change their expression and affect seed germination. Studies showed the role and the versatility of mitochondria structures during seed germination. At dry seed stage there are promitochondria, which are larger, isodiametric structures that will eventually develop into mitochondria upon seed imbibition. Later, upon germination, mitochondria undergoes variations in morphology along with cell differentiation and cell division in the course of early root development [118,119]. The bioenergetic reactivation of mitochondria is immediate observed upon seed imbibition and the reactivation of mitochondrial dynamics occurs after transfer to germination conditions [120]. Besides, as mitochondria are the main source of reactive oxygen species (ROS) production, they are strongly exposed to oxidative damage [121], and the continuous heat stress could affect them greatly, also in an epigenetic manner, as we observed in our work. Further studies on these genes could clarify their role on heat stress resistance in seeds.

Taken together, our results showed a partial control of heat stress response on seeds by the demethylation of germination-related genes. This is a finding of great importance because seed germination and plantlet establishment are the base of global food production, and ensuring food security in the climate change era is one of the major challenges in the coming decades. Plant breeding strategies focused on selecting varieties with increased resistance to abiotic stresses, like heat stress, will play a central role in confronting this challenge.

## Supporting information

STable1

STable2

SFig 1

SFig 2

SFig 3

## Supplementary material

**Figure S1. Col-0 plant phenotype under heat stress conditions.** (A) Photos of Col-0 plants showing increased biomass related to increase in temperature. (B) Photos of Col-0 plants showing decreased silique number related to the increase in temperature. (C) Average silique number per branch at 21°C, 23°C, 25°C, 27°C and 29°C average temperatures. (D) Average silique length at 21°C, 23°C, 25°C, 27°C and 29°C average temperatures. Standard deviations are indicated in both C and D graphs.

**Figure S2. Microscopy of Col-0 embryo development under heat stress.** Embryo stages showed are globular, transition, heart, torpedo, walking stick and mature at 21°C, 23°C, 25°C and 27°C average temperature.

**Figure S3. Gene set enrichment analysis (GSEA) of specific DEGs in Col-0 seeds under severe heat stress.** (A) Enriched GO terms from up-regulated differentially expressed genes. (B) Enriched GO terms from down-regulated differentially expressed genes. The size of the dot represents the gene count. The totality of differentially up- or down-regulated genes specific to each stage was used to perform a hypergeometric test, and the p-values from the tests were converted to false discovery rate (FDR)-corrected p-value as shown in colours, the red colour being more significant than blue colour. H= heart, B= bent, M= mature and C= common to all stages.

**Supplementary Table 1:** Deseq2 summary data from RNA-seq with Col-0 seeds during development under different ranges of heat stress.

**Supplementary Table 2:** Deseq2 summary data from RNA-seq with *ros1-4* and Col-0 seeds at maturation and germination stages.

**Supplementary Data:** RNA-seq and methylome data are available at GSE167245 (https://www.ncbi.nlm.nih.gov/geo/query/acc.cgi?acc=GSE167245).

## Acknowledgements

The authors sincerely thank all SEED team at INRAE, IRHS from Angers, more specifically Joseph Ly Vu and Benoit Ly Vu for advices regarding plant care and physiological assays. The authors gratefully acknowledge the technical platforms ANAN and iMAC of the SFR 4207 QUASAV.

## Authors contribution

Conceptualization, JM, DW and JV; Methodology, JM, DW, WX, JV; Validation, JM, JV; Formal Analysis, JM, JV, WX and DW; Investigation, JM, DW, WX and JV; Data Curation, JM and JV; Writing – Original Draft Preparation, JM; Writing – Review & Editing, JV, DW, WX and JM; Visualization, JM and JV; Supervision, JV; Project Administration, JV; Funding Acquisition, JV.

