## Supplementary figures and images for "Regulation of DNA (de)methylation positively impacts seed germination during seed development under heat stress"

### SFig 1

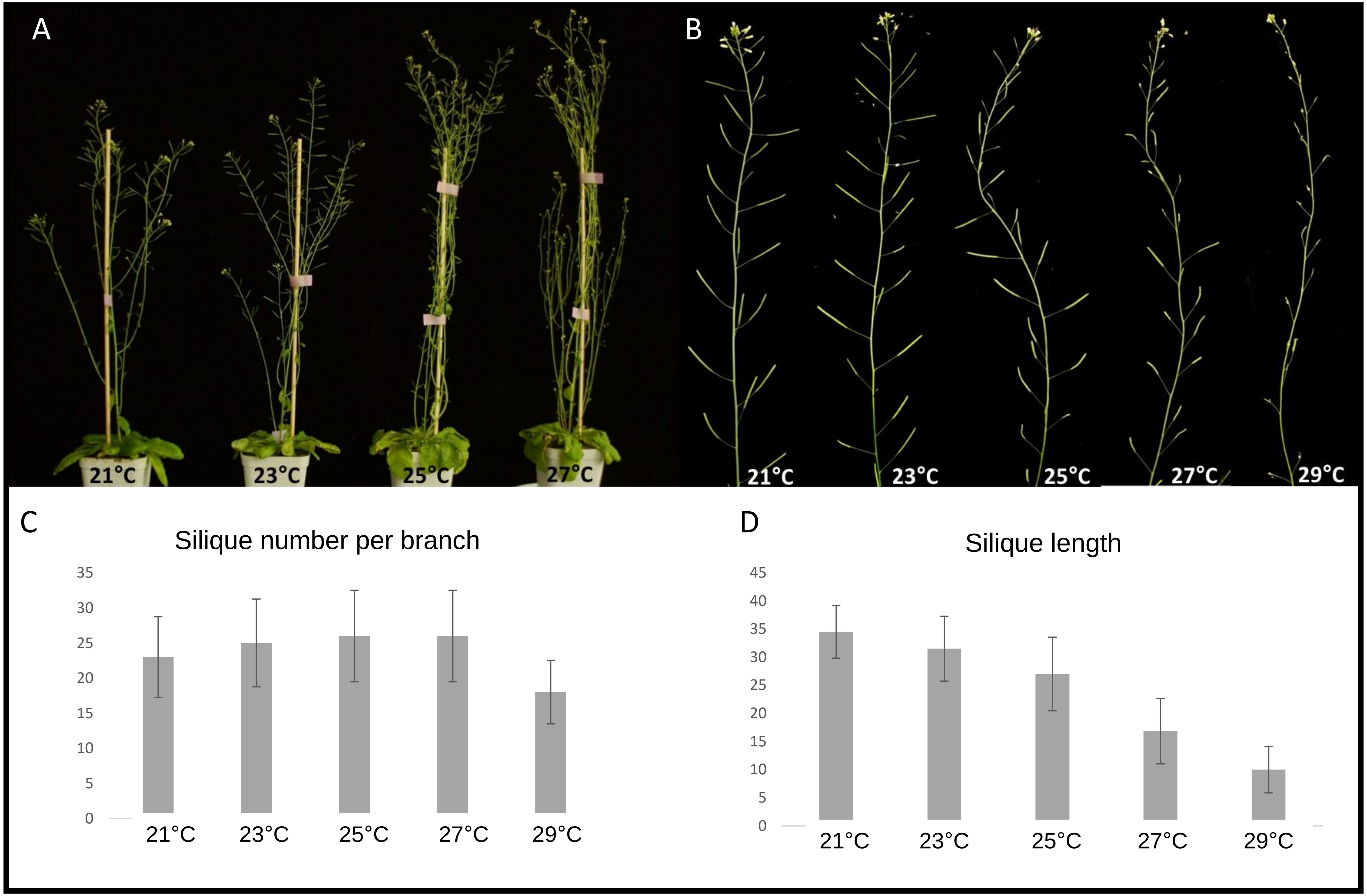

### SFig 2

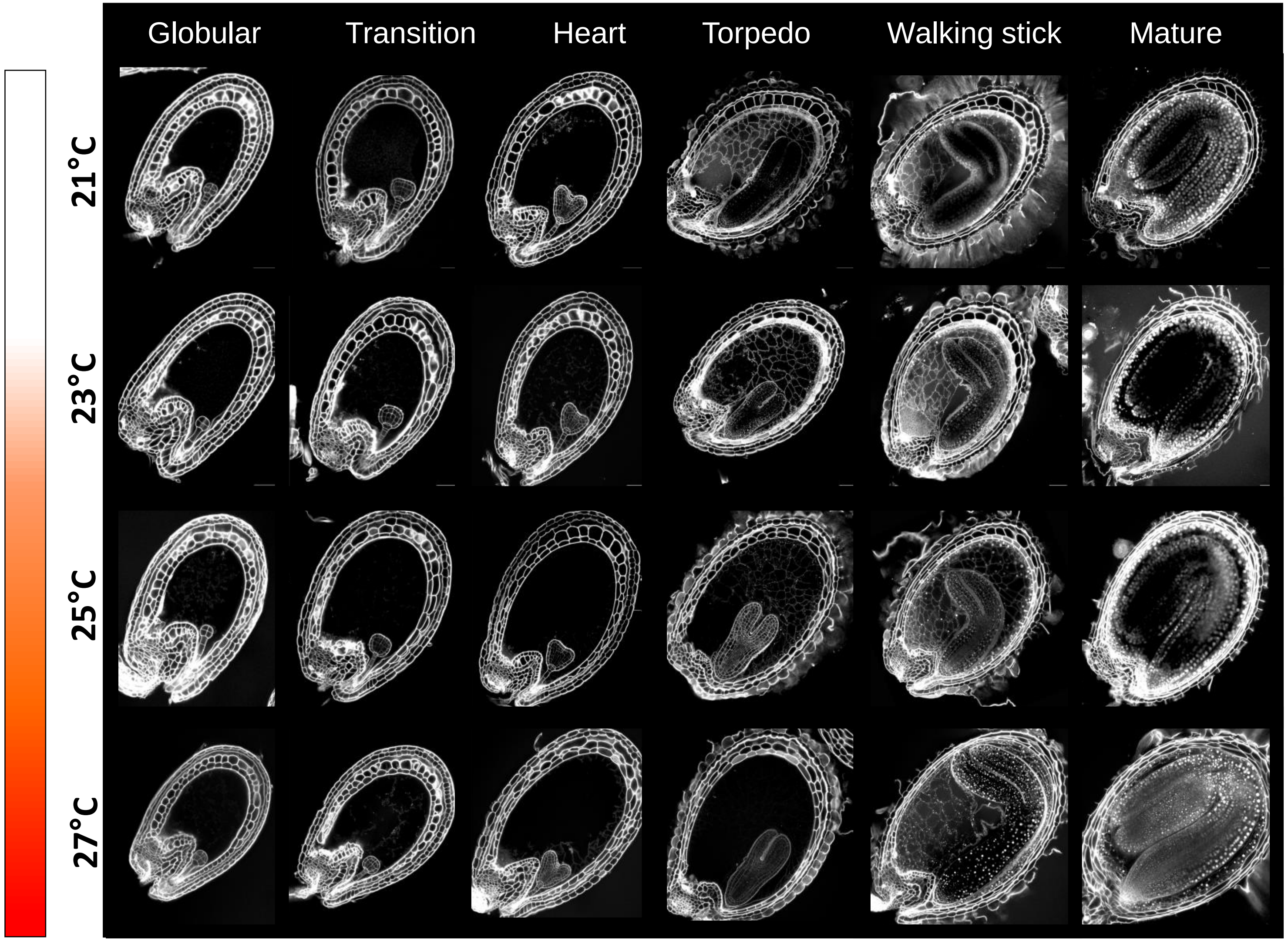

### SFig 3

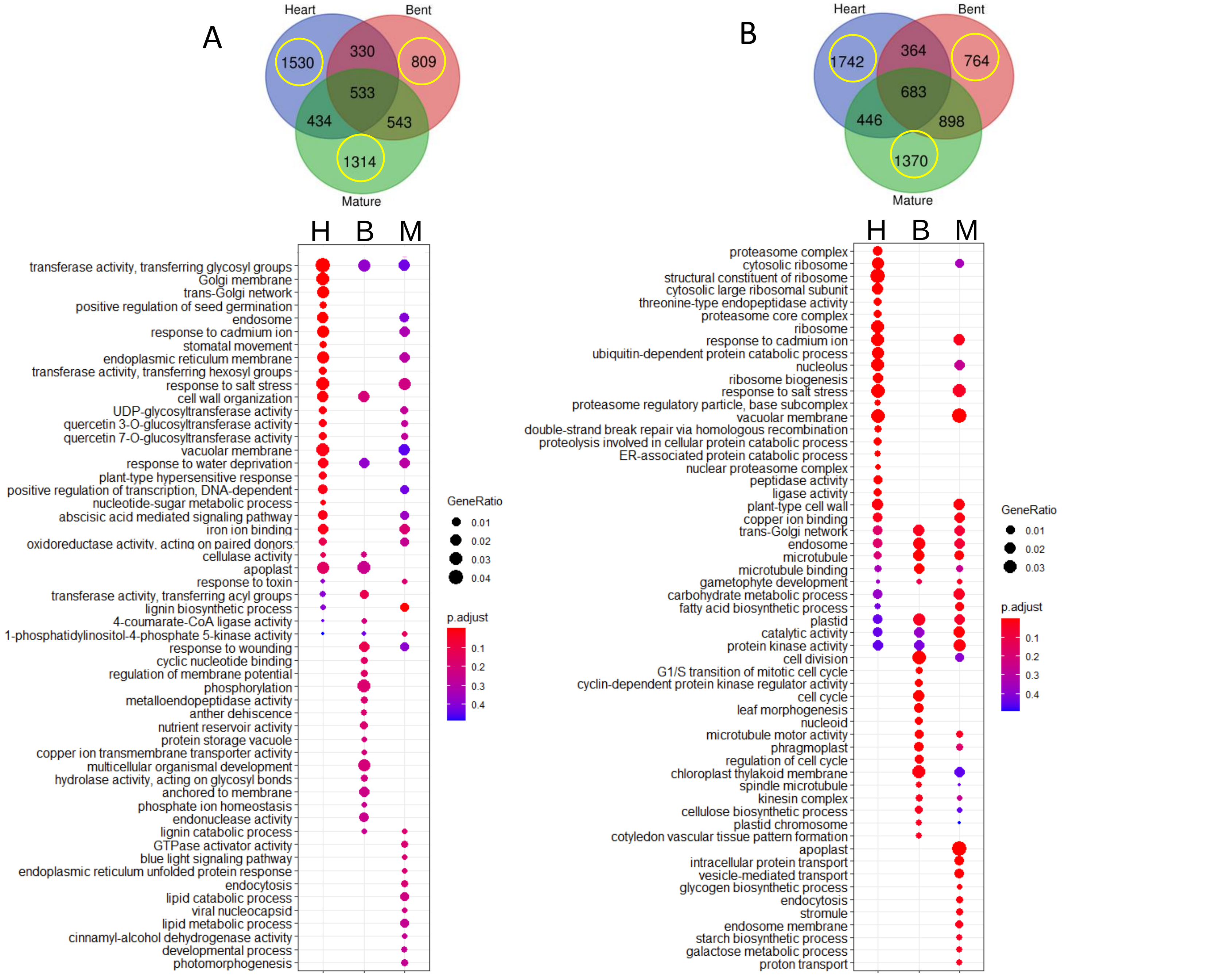
